# OmniBind: Proteome-Wide Promiscuity Predictions for Early-Stage Drug Screening

**DOI:** 10.64898/2026.03.17.712433

**Authors:** Josef Hanke, Sebastian Pujalte Ojeda, Ruoh Wen Cheong, Logan M. Glasstetter, Ellis Baker, Hilbert Lam, Michaela Brezinova, Adelie Louet, Shengyu Zhang, Michele Vendruscolo

## Abstract

Off-target binding remains a leading cause of drug attrition, yet no method exists for rapidly quantifying small-molecule promiscuity across the human proteome. Here, we define promiscuity as the mean predicted binding affinity over 15,405 human proteins and derive a specificity score combining target affinity with this proteome-wide distribution. To make this assessment tractable at scale, we introduce OmniBind, a message-passing neural network that predicts promiscuity directly from a SMILES string at about a thousand compounds per second, several orders of magnitude faster than proteome-wide profiling. OmniBind promiscuity scores correlate with experimental binding data near assay reproducibility limits. Ranking candidates by specificity rather than affinity alone improves enrichment of approved drugs across all thresholds tested, an advantage robust to the choice of affinity predictor. OmniBind fills an unoccupied niche in the early-stage screening landscape as a fast, proteome-scale complement to traditional safety panels, with accuracy that will scale as the underlying affinity predictors continue to improve.

## Introduction

Drug discovery remains one of the most expensive and failure-prone enterprises in the biomedical sciences. Global biopharmaceutical R&D investment exceeds $276 billion annually^1^, yet the cost per approved drug continues to rise^2^, and only about 10% of compounds entering phase I clinical trials ultimately reach the market^3,4^. Among the leading causes of attrition are insufficient efficacy and off-target toxicity, which can in some cases both be influenced by non-selective target engagement. A drug that binds promiscuously across the proteome will have less free compound available to engage its intended target, reducing efficacy^5^, while simultaneously increasing the probability of triggering adverse effects through unintended interactions^6^. On average, approved small-molecule drugs are estimated to bind several off-target proteins^7-9^, and the true number is likely higher given the limited scope of typical screening methods.

Despite the centrality of this problem, promiscuity and specificity remain imprecisely defined concepts with no consensus quantitative framework^10-12^. In current practice, candidate compounds are screened for off-target liability against small panels of proteins selected on the basis of known toxicity associations or structural similarity to the intended target^6,13-16^, an approach intended to improve clinical attrition rates^17^. While informative, these panels are inherently incomplete, as they cannot detect binding to structurally dissimilar off-targets or to proteins with as-yet-uncharacterised toxicities^17-19^. Notably, no computational or experimental screening panel includes disordered proteins, which constitute over one third of human protein-coding genes^20,21^ and include pathologically important targets such as amyloid-β^22,23^ and α-synuclein^24,25^. A recent industry-wide analysis found that although more recently approved drugs show reduced promiscuity against standard panels such as the Bowes-44^6,17^, neither safety-related attrition rates nor overall clinical trial success rates have meaningfully improved^26-28^, suggesting that current screening strategies remain insufficient.

Computational approaches have begun to address this gap. Structure-based methods such as molecular docking evaluate binding in a one-to-one paradigm in which a given ligand binds a target protein^29,30^, and are restricted to targets with well-resolved binding pockets, estimated to cover roughly 3,000 proteins in the druggable genome^31^ or up to one third of the proteome when computational pocket prediction is included^18,32^. Machine-learning models for drug-target affinity prediction have improved accuracy and throughput considerably^33-35^, and machine learning has been combined with limited-proteolysis chemoproteomics to identify drug targets and binding sites across complex proteomes^36^, yet they are deployed in a similar paradigm in which the affinity to the intended target is optimised first, and off-target liability is assessed downstream, against restricted panels^17,37-39^. A recent study docked 7,582 drugs against 19,135 AlphaFold-predicted human protein structures and used the resulting binding profiles to predict therapeutic indications and side effects, but did not define a continuous promiscuity metric or combine it with target affinity for candidate ranking^40^. There is thus a need for computational methods to define a promiscuity score based on proteome-wide binding behaviour and that can be combined with target affinity predictions to yield a unified specificity metric suitable for integration into screening or de novo design pipelines.

Here we concentrate on a shift from this one-to-one paradigm to a one-to-all model of molecular recognition. We define promiscuity as the mean predicted binding affinity of a small molecule across a sizeable fraction of the human proteome (15,405 proteins), computed using Ligand-Transformer, a deep-learning model that predicts binding affinity from protein sequence and molecular structure and that, unlike docking-based approaches, can score interactions with disordered proteins^35^.

From this proteome-wide binding distribution, we derive a specificity score as a Z-score that quantifies how far the affinity of a compound for a given target deviates from its average affinity across the proteome. We show that this proteome-wide promiscuity metric correlates strongly (Spearman ρ > 0.96) with abundance-weighted, tissue-specific formulations across blood plasma, liver, brain, and whole-body contexts, supporting its use as a general-purpose measure that does not require tissue-specific parameterisation.

Because computing a full proteome binding profile for every candidate molecule is expensive, we introduce OmniBind (Promiscuity Predictor), a message-passing neural network (MPNN)^41^ built on the Chemprop framework^42,43^, which predicts promiscuity and its associated standard deviation directly from the SMILES representation of a compound^44^. OmniBind achieves a mean squared error of 0.016 pK_D_ on held-out test data and processes about a thousand molecules per second, which is roughly five orders of magnitude faster than computing full proteome profiles with Ligand-Transformer^35^ and eight orders of magnitude faster than other approaches such as Boltz-2^34^.

We validate OmniBind predictions against experimental binding data from multiple independent databases, showing that computational-experimental agreement approaches the reproducibility limits of the experimental measurements themselves. We further demonstrate that OmniBind promiscuity scores correlate negatively with clinical maximum unbound plasma concentrations (C_max_), obtained from independent pharmacokinetic databases^45,46^, at a level approaching the inter-database reproducibility of C_max_ measurements themselves. This correlation is consistent with the free drug hypothesis^5^ and persists after controlling for molecular weight^47^, providing pharmacokinetic evidence that the metric captures biologically meaningful variation beyond simple physicochemical properties. When used to rank FDA-approved drugs^48^ among random decoy compounds from the TargetMol database (https://www.targetmol.com/), specificity scoring consistently outperforms affinity-alone ranking across all enrichment thresholds tested, as measured by both BEDROC^49^ and enrichment factor metrics, an improvement that is robust to the choice of underlying affinity predictor, as confirmed by independent analysis with Boltz-2^34^ and on the Eurofins SafetyScreen panel^46^ (https://www.eurofinsdiscovery.com/solution/safety-panels). To make proteome-wide promiscuity assessment broadly accessible, we report the OmniBind web server, which allows researchers to obtain promiscuity and specificity predictions within seconds from a standard web browser.

## Results

### A proteome-wide promiscuity metric

We used Ligand-Transformer to predict binding affinities between ∼61,000 small molecules and 15,405 human proteins, yielding a complete proteome-wide binding profile for each compound (**Figure S1**). From these profiles, we computed two candidate promiscuity metrics: a simple unweighted mean affinity across the proteome, and a tissue-specific formulation that weights each protein by its measured abundance in a given tissue (Methods). We evaluated the tissue-specific metric across several drug-relevant contexts (blood plasma, liver, brain, the HEK293 cell line, and a whole-body average) and found that all formulations correlate almost perfectly with the unweighted mean (Spearman ρ > 0.96 in all cases; **Figure 1**). This consistency indicates that promiscuity depends more on the global binding tendency of a molecule than on the expression profile of any particular tissue. We therefore adopted the unweighted mean affinity as our promiscuity metric for all subsequent analyses, as it requires no tissue-specific parameterisation and generalises across biological contexts.

**Figure 1.**
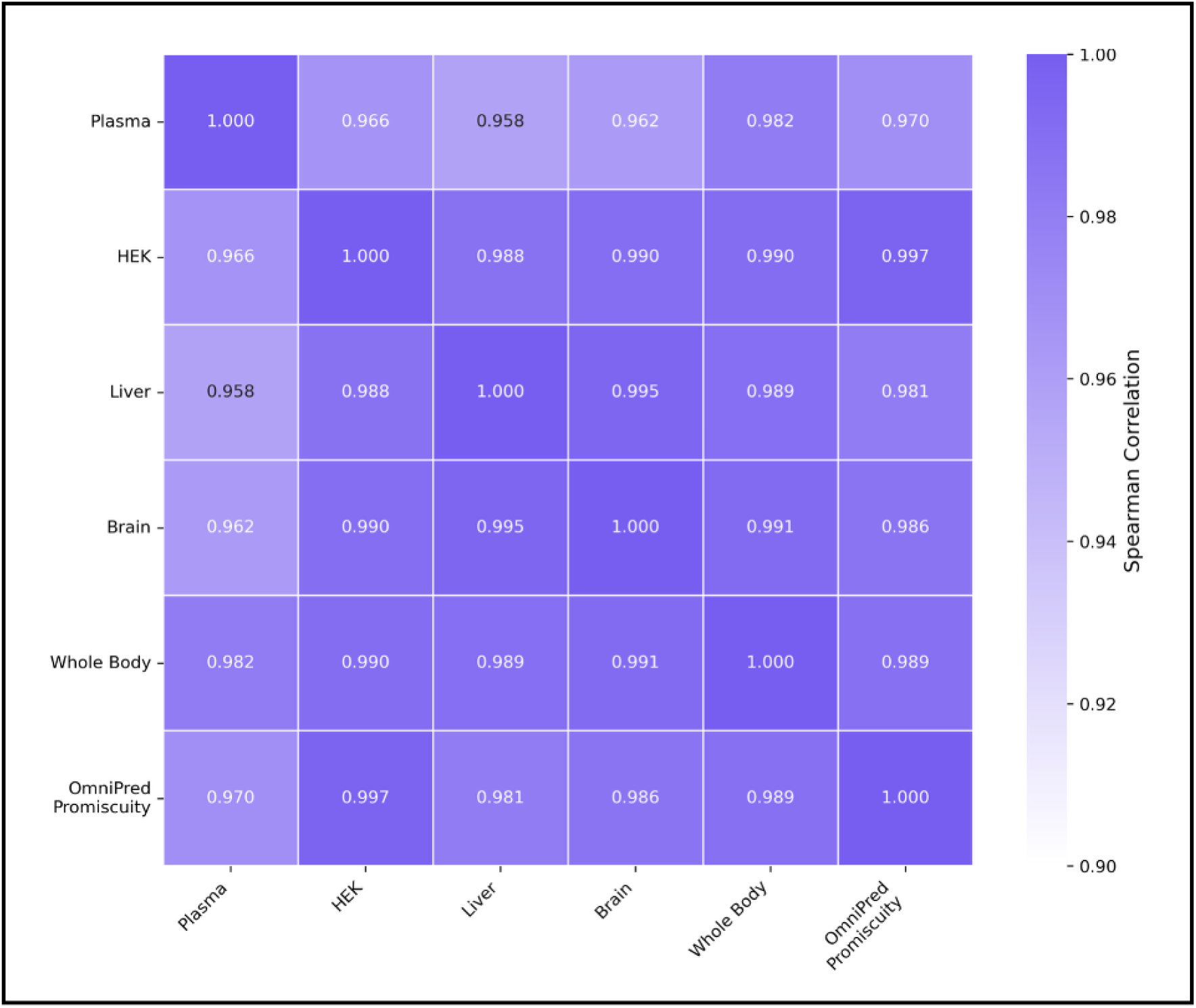
Correlation between tissue-specific and proteome-wide promiscuity metrics. Heatmap showing the Spearman correlation between tissue-weighted promiscuity scores and the unweighted mean binding affinity across the human proteome (15,405 proteins). Tissue-specific formulations weight each protein by its measured abundance in the corresponding biological context (blood plasma, liver, brain, HEK293 cells, and whole-body average). All tissue-weighted metrics correlate strongly with the unweighted mean (Spearman ρ > 0.96), indicating that promiscuity is largely determined by the global binding tendency of a molecule rather than by tissue-specific protein expression profiles.

### OmniBind: a fast promiscuity predictor

Computing a full proteome binding profile via Ligand-Transformer is accurate but expensive, requiring ∼15,000 individual affinity inferences per molecule. To make promiscuity assessment practical at scale, we trained OmniBind, an MPNN that predicts the promiscuity score of a molecule and the standard deviation of its binding profile directly from its SMILES representation (**Figure S1**). Training proceeded in two stages: first, a random forest regressor learned to estimate promiscuity from a subsample of only the 173 most representative protein affinities (∼1% of the proteome). These 173 proteins were selected using an importance metric from a random forest regressor trained on the whole proteome. This intermediate model achieved an mean square error (MSE) of 0.03 against the complete proteome promiscuity (CPP) scores (Methods), and was used to generate 200,935 additional training labels at a fraction of the cost of full proteome inference; second, these augmented labels, together with 61,677 complete proteome promiscuity scores, were used to train the final MPNN model (Methods).

OmniBind was evaluated on 1,000 held-out TargetMol molecules with known complete proteome promiscuity scores. It predicted both the mean and standard deviation of binding affinities with mean squared errors of 0.016 and 0.001, respectively (**Figure S2**). Inference time scales linearly with the number of molecules, processing about a thousand molecules per second. This is roughly five orders of magnitude faster than equivalent Ligand-Transformer-based proteome profiling.

We additionally confirmed that Ligand-Transformer and Boltz-2 affinity predictions for drug– target pairs correlate strongly (|r| ≥ 0.6, p ≤ 0.001; **Figure S3**), although OmniBind is about eight orders of magnitude faster than Boltz-2 (**Figure S4**), supporting the validity of using Ligand-Transformer as the primary affinity predictor in this study.

### Validation of promiscuity scores

Validating a proteome-wide promiscuity metric against experimental data is inherently challenging, as no assay directly measures the mean binding affinity of a compound across the entire proteome. As a proxy, we assembled binding affinity data for small molecule-protein pairs from three independent databases (ChEMBL^50^, the PDSP Ki database^51^, and the Safety Pharmacology Database^45^) and derived four distinct experimental promiscuity estimates (Methods) (**Figure 2**). These proxy scores correlate only moderately with one another (r=0.25-0.49), reflecting the limited and variable protein coverage of each dataset (typically fewer than ten proteins per molecule) (**Figure 2**). Despite this, OmniBind promiscuity scores correlate with the ChEMBL Ki and PDSP Ki proxies at levels approaching the inter-database agreement, indicating that computational and experimental estimates of promiscuity converge at a limit set by the incompleteness of existing experimental data rather than by the computational method.

**Figure 2.**
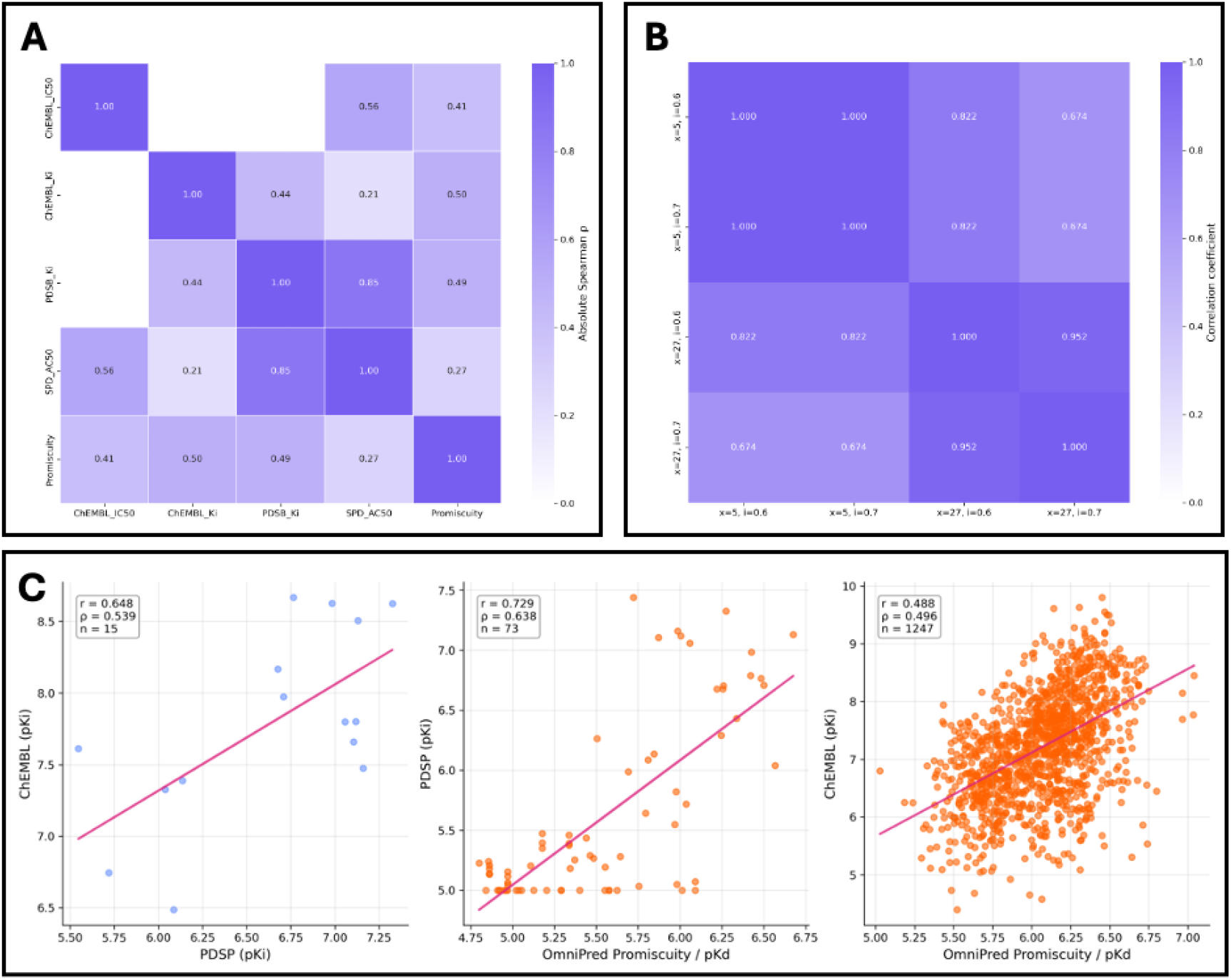
Validation of proteome-wide promiscuity scores against experimental binding datasets. **(a**,**b)** Experimental proxy measures of compound promiscuity derived from binding affinity data in ChEMBL, the PDSP Ki database, and the Safety Pharmacology Database. For each compound, promiscuity proxies were calculated from the available set of measured protein-ligand affinities (Methods). **(c)** Pairwise correlations between the different experimental promiscuity proxies, showing moderate agreement (r = 0.25-0.49) due to the limited and heterogeneous protein coverage of each dataset. Despite this variability, OmniBind promiscuity scores correlate with experimental proxies at levels approaching the inter-database agreement, indicating that computational and experimental estimates of promiscuity converge within the limits imposed by incomplete experimental coverage.

### Correlation with maximum unbound plasma concentration

A pharmacokinetic parameter of particular clinical relevance is C_max_, the maximum unbound drug concentration achieved in plasma. A promiscuous molecule that binds extensively across the proteome should, by the free drug hypothesis, exhibit a lower C_max_ at a given dose. We tested this prediction using C_max_ measurements from two independent databases (SPD^45^ and PKDB^46^), filtered for healthy adults in blood plasma (see Methods). The experimental C_max_ values correlate with each other with r=0.55, establishing an upper bound on the reproducibility of this measurement. OmniBind promiscuity scores correlate negatively with C_max_ from both databases (r=-0.47 and r=-0.45, respectively), approaching this inter-database reproducibility limit (**Figure 3**, left panels).

**Figure 3.**
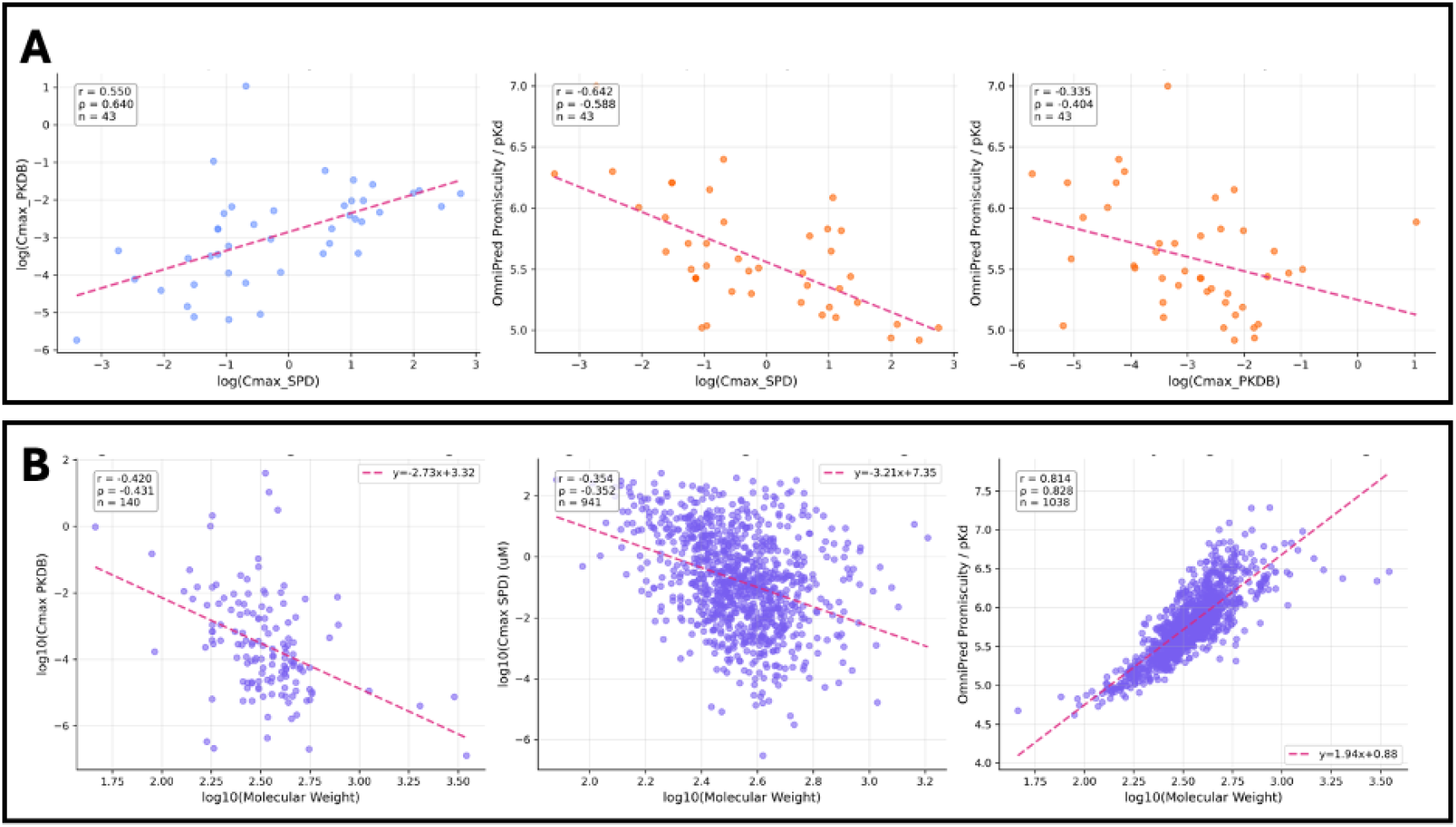
Promiscuity scores correlate with maximum unbound plasma concentration (C_max_) beyond molecular weight alone. Left panels, correlations between OmniBind promiscuity scores and experimentally measured maximum unbound plasma concentrations (C_max_) from the SPD and PK-DB databases for compounds with measurements in healthy adults in blood plasma. Right panels, the corresponding correlations between molecular weight and the same Cmax values. Each point represents one compound; lines indicate best-fit linear regressions. Experimental C_max_ values from the two databases correlate with each other with r=0.55, while OmniBind promiscuity scores correlate negatively with C_max_ in both datasets (r=-0.47 and r=-0.45), exceeding or matching the corresponding correlations with molecular weight alone (r=-0.35 and r=-0.42). These results indicate that promiscuity captures pharmacokinetically relevant variation beyond simple size-related physicochemical effects.

Because promiscuity correlates strongly with molecular weight (Pearson r=0.86), we verified that the C_max_ relationship is not simply a molecular weight effect. Correlating the same C_max_ values with molecular weight alone yields substantially weaker associations (r=-0.35 and r=-0.42; **Figure 3**, right panels), confirming that promiscuity captures pharmacokinetically relevant variation beyond simple physicochemical properties. Importantly, OmniBind was trained entirely independently of C_max_ data, making this correlation an external validation of the biological meaning of the metric.

### Promiscuity and specificity of approved drugs

FDA-approved drugs exhibit significantly lower promiscuity than random small molecules (mean 5.75 pK_D_ versus 5.88 pK_D_; p < 0.001, Student’s t-test; **Figure 4**), consistent with the expectation that the drug development process selects against compounds with extensive off-target binding.

**Figure 4.**
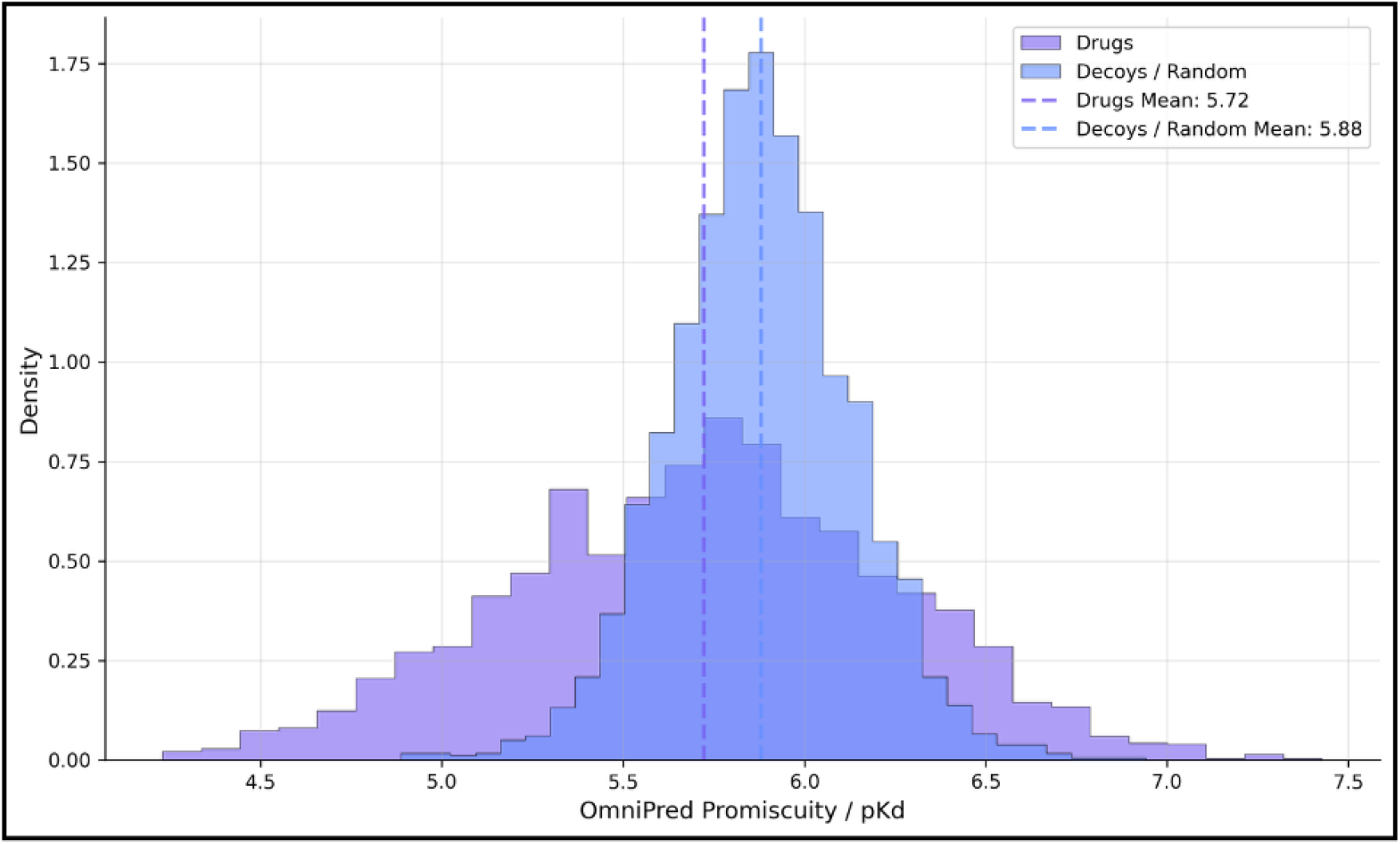
FDA-approved drugs exhibit lower predicted promiscuity than random small molecules. Overlaid histograms showing the distribution of OmniBind promiscuity scores for 2,663 FDA-approved drugs and 2,663 randomly selected compounds from the TargetMol database. Promiscuity is defined as the mean predicted binding affinity across 15,405 human proteins. FDA-approved drugs are shifted toward lower promiscuity values than random compounds (mean 5.75 pK_D_ versus 5.88 pK_D_; Student’s t-test, p < 0.001), consistent with the idea that drug development selects against compounds with extensive off-target binding across the proteome.

We next examined the relationship between promiscuity and target engagement. For each approved drug, we predicted its affinity to all annotated DrugBank^48^ targets using Ligand-Transformer and selected the highest-affinity target (affinity to best target, ABT). ABT correlates positively with promiscuity (r = 0.60, p < 0.001; **Figure S5**), indicating that drugs with high target affinity also tend to be more promiscuous, a finding that underscores the need to evaluate both properties jointly rather than optimising affinity in isolation.

To this end, we combined the ABT of each drug with its OmniBind promiscuity score to compute a specificity score (a Z-score describing the affinity to the target relative to the distribution of affinities across the proteome; Methods). **Figure S6** shows the distribution of specificity scores across drug categories. Allergy medications, psychiatric drugs, and sedatives exhibit the highest mean specificities (3.10, 2.76, and 2.76, respectively), whereas diagnostics, antiseptics, and antibiotics have the lowest (0.25, 0.46, and 0.58, respectively), a pattern consistent with the pharmacological requirements of each class.

### Specificity improves drug enrichment

To quantify the practical value of the specificity metric, we evaluated its ability to distinguish approved drugs from random compounds. For each drug target, we ranked the corresponding drug alongside 1,000 randomly selected TargetMol decoys, using either affinity alone or our combined specificity score. After excluding drug categories for which specificity is pharmacologically less relevant (antiseptics, dermatologicals, diagnostics, nutritional supplements, ophthalmics), ranking by specificity consistently placed approved drugs higher than ranking by affinity alone. This improvement was significant across all BEDROC α values and enrichment factor thresholds tested (Table 1). For example, the top-1% enrichment factor rose from 11.26 (affinity) to 12.49 (specificity), and BEDROC at α = 10 increased from 0.251 to 0.308.

**Table 1.** Specificity-based ranking improves enrichment of approved drugs over affinity alone. Enrichment of FDA-approved drugs among 1,000 randomly sampled TargetMol decoy compounds when ranked by the combined specificity score versus predicted target affinity (pK_D_) alone. BEDROC scores are reported at three α values controlling the weight given to early recognition, and enrichment factors are reported at four rank thresholds. Higher values indicate better separation of approved drugs from decoys. Drug categories for which target specificity is pharmacologically less relevant (antiseptics, dermatologicals, diagnostics, nutritional supplements, and ophthalmics) were excluded.

| Method | BEDROC |  |  | Enrichment Factor |  |  |  |
| --- | --- | --- | --- | --- | --- | --- | --- |
| | $\alpha = 10$ | $\alpha = 20$ | $\alpha = 80$ | EF 1% | EF 5% | EF 10% | EF 20% |
| Affinity | 0.251 | 0.188 | 0.105 | 11.26 | 4.32 | 2.85 | 1.85 |
| Specificity | **0.308** | **0.223** | **0.113** | **12.49** | **5.14** | **3.40** | **2.35** |

To confirm that this result is robust, we repeated the analysis using 5-fold cross-validated decoy splits (n = 192 decoys per fold). The specificity advantage was reproduced in every fold, with the improvement reaching statistical significance across all enrichment metrics (one-sided paired Wilcoxon, p < 0.05; **Table S1**). The cumulative ranking curves (**Figure 5**) illustrate this visually: specificity-ranked drugs (solid lines) consistently dominate affinity-ranked drugs (dashed lines) at every rank threshold, with the median rank improving from 45 ± 0 (affinity) to 29 ± 1 (specificity) when using Ligand-Transformer.

**Figure 5.**
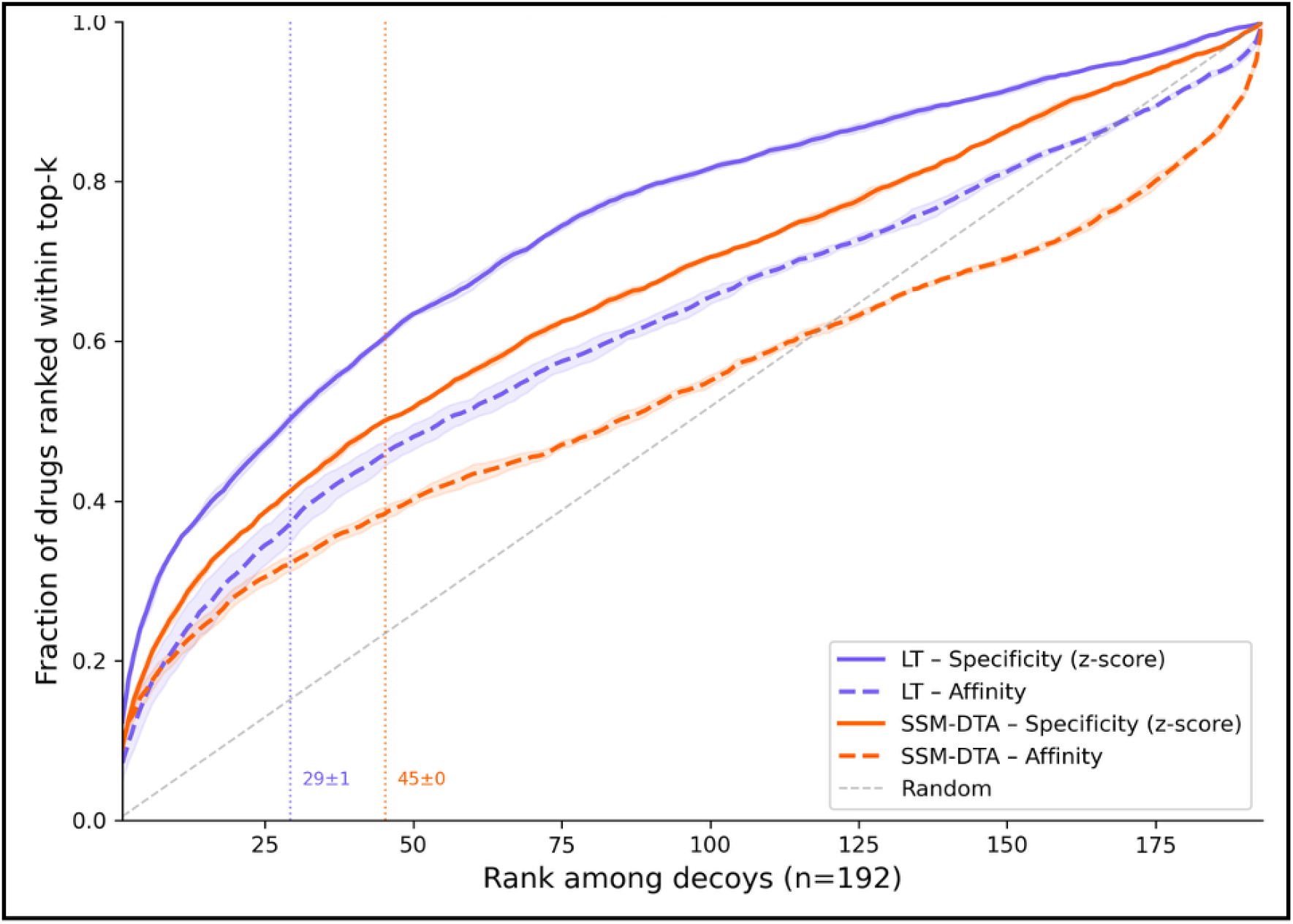
Specificity-based ranking consistently outperforms affinity-only ranking across affinity predictors. Cumulative ranking curves showing the proportion of FDA-approved drugs recovered as a function of rank position when drugs are ranked among random TargetMol decoys using either affinity alone (dashed lines) or the combined specificity score incorporating proteome-wide promiscuity (solid lines). Results are shown for both Ligand-Transformer (LT) and SSM-DTA as the underlying affinity predictor, evaluated across five-fold cross-validated decoy splits (n = 192 decoys per fold). Specificity-ranked drugs consistently dominate affinity-ranked drugs at every rank threshold across both predictors, with the median rank improving from 45 ± 0 to 29 ± 1 when using Ligand-Transformer. LT-based scoring outperforms SSM-DTA-based scoring in both the affinity and specificity conditions, consistent with the higher reported accuracy of Ligand-Transformer. The advantage of incorporating promiscuity is not dependent on the particular affinity predictor used.

### Generalisability across affinity predictors

A key question is whether the specificity improvement depends on the particular affinity predictor used. We addressed this in two ways.

First, we replaced Ligand-Transformer with SSM-DTA^52^, an independent drug-target affinity model, and repeated the 5-fold enrichment analysis. The specificity advantage persisted: SSM-DTA specificity outperformed SSM-DTA affinity across all metrics, with improvements comparable to those observed with Ligand-Transformer (**Table S1 and Figure 5**). Notably, LT-based scoring outperformed SSM-DTA-based scoring in both the affinity and specificity conditions, consistent with the higher reported accuracy of Ligand-Transformer on binding affinity benchmarks.

Second, we tested whether promiscuity, as captured by our metric derived from Ligand-Transformer, correlates with binding profiles predicted by a structurally distinct method. We used Boltz-2 to predict the affinity of 63 approved drugs to each of the 53 proteins in the Eurofins SafetyScreen panel and computed the mean Boltz-2 affinity score for each drug across the panel. These Boltz-2 panel scores correlate strongly with Ligand-Transformer promiscuity (r = −0.74, p < 10^−11^; **Figure S7**, left panel). Furthermore, drugs classified as highly promiscuous by Ligand-Transformer (mean pK_D_ ≥ 5.85) appear significantly more often among the strongest 10% of Boltz-2 binders across the 53 Eurofins proteins than do low-promiscuity drugs (Mann-Whitney p = 7.0 × 10^−6^; **Figure S7**, right panel).

## Discussion

We have introduced a quantitative framework for small-molecule promiscuity and specificity grounded in proteome-wide binding predictions, and illustrated how these metrics have both biological meaning and practical value for drug screening. Three principal findings emerge.

First, a simple, unweighted mean of predicted binding affinities across about 15,000 human proteins serves as a robust measure of promiscuity. Despite the considerable biological heterogeneity across tissues arising from differences in protein expression, drug delivery routes, and individual physiology, the unweighted promiscuity score correlates almost perfectly (Spearman ρ > 0.96) with abundance-weighted, tissue-specific formulations for blood plasma, liver, brain, and whole-body contexts (**Figure 1**). This consistency is notable because it means the metric does not require tissue-specific parameterisation, which would reduce generalisability and increase computational cost without meaningfully improving the characterisation of the global binding tendency of a molecule. The practical implication is that a single number, the OmniBind promiscuity score (OPS), can serve as a general-purpose descriptor of off-target liability across biological contexts.

Second, the OPS captures biologically meaningful variation beyond what molecular weight alone can explain. Promiscuity correlates strongly with molecular weight (Pearson r = 0.86), as expected given that larger molecules present more functional groups for interaction. However, when we correlate OPS with experimentally measured maximum unbound plasma concentrations (C_max_) from two independent pharmacokinetic databases, the resulting correlations (r=-0.47 and r=-0.45) substantially exceed those between C_max_ and molecular weight alone (r=-0.35 and r=-0.42), and approach the reproducibility limit of C_max_ measurements across databases (r = 0.55). Crucially, OPS was trained entirely independently of C_max_ data. The observed negative correlation between OPS and C_max_ is directionally consistent with the free drug hypothesis, although C_max_ reflects the combined influence of absorption, distribution, metabolism, and clearance rather than protein binding in isolation. The similarity in magnitude between the OPS-C_max_ and inter-database C_max_ correlations sets a practical ceiling on what can be resolved from these data, but does not establish that the computational metric has reached an intrinsic accuracy limit, as controlling for additional physicochemical covariates may alter the residual correlation. We have obtained complementary external validation from experimental binding databases. The correlation between OPS and proxy promiscuity scores derived from ChEMBL and PDSP Ki data approaches the correlation between these experimental proxies themselves, indicating that computational and experimental estimates of promiscuity agree at a level limited primarily by the incompleteness of existing experimental data. Orthogonal evidence comes from proteome-wide expression profiling, where compound-induced changes in protein abundance across near-complete proteomes^53^ provide an independent readout of the breadth of the cellular effects of a molecule that could, in future work, be correlated with OmniBind promiscuity scores.

Third, integrating promiscuity with target affinity into a specificity score significantly improves the identification of approved drugs within random compound libraries. Ranking by specificity rather than affinity alone improves enrichment at every threshold tested. For example, BEDROC at α = 10 increases from 0.251 to 0.308, and the top-1% enrichment factor rises from 11.26 to 12.49 (Table 1). This improvement is not an artefact of the affinity predictor used: it is reproduced when Ligand Transformer predictions are replaced with Boltz-2 affinity scores, and when evaluated against the industry-standard Eurofins SafetyScreen panel, where promiscuity correlates strongly with Boltz-2 binding profiles (r=−0.74, p<10^™11^). The consistency across affinity predictors and evaluation contexts supports the conclusion that the specificity improvement indicates that approved drugs are selected not only for strong target engagement but also, implicitly, for low off-target liability.

The analysis of drug categories (**Figure S6**) provides additional face validity. Allergy medications, psychiatric drugs, and sedatives (categories that require high central nervous system penetration and thus benefit from low plasma protein binding) exhibit the highest specificity scores. Conversely, topical agents (antiseptics, dermatological drugs) and compounds not designed to bind specific protein targets (diagnostics, nutritional supplements) have the lowest scores, consistent with the expectation that specificity is less relevant to their therapeutic function. We excluded these latter categories from the enrichment analysis to avoid biasing the evaluation, but their behaviour is itself informative, as it confirms that the specificity score tracks the pharmacological logic of different drug classes.

Several limitations should be noted. The promiscuity score is a coarse-grained, proteome-wide summary and is not designed to resolve binding to individual off-target proteins. Adverse effects are most often driven by potent binding to one or a small number of specific off-targets, and identifying these requires the kind of protein-level resolution that panel-based screening provides. OmniBind is therefore best understood as a complement to, rather than a replacement for, existing experimental and computational safety screens: it offers a global overview of off-target liability that can inform early prioritisation decisions, while traditional panels remain essential for characterising specific safety-relevant interactions. A further limitation is the exclusion of the longest 20% of human proteins from the reference proteome, imposed by the quadratic scaling of Ligand-Transformer inference time with protein length. Although the remaining 15,000 proteins span the majority of human protein classes, and the high tissue-correlation results suggest the promiscuity score is robust to this exclusion, some potential off-target interactions with very large proteins may be missed. Finally, the current framework considers only protein off-targets; binding to nucleic acids, lipids, and other biomolecules is not captured.

OmniBind can process about a thousand compounds per second, roughly five orders of magnitude faster than Ligand-Transformer and eight orders faster than Boltz-2 for equivalent proteome-wide profiling. This computational efficiency makes it practical to incorporate promiscuity assessment into high-throughput and generative drug design workflows via OmniBind. To make this capability broadly accessible, we report the OmniBind web server, which accepts individual or batch SMILES submissions and returns promiscuity scores, standard deviations, and optional specificity scores within seconds.

We anticipate a near-term application in the integration of specificity scoring as an optimisation objective in generative molecular design frameworks such as SynFlowNet^54^, where OmniBind can serve as a lightweight oracle guiding exploration of chemical space toward molecules that are not only potent against their intended target but also selective across the proteome. Looking further ahead, extending the framework to include binding predictions for nucleic acids and safety-relevant protein panels would yield a more comprehensive picture of molecular selectivity and could support the development of unified, multi-objective scoring functions for early-stage drug design.

## Methods

### Promiscuity definitions

We considered two candidate definitions of promiscuity. The first is a tissue-specific formulation derived from equilibrium binding theory. Given a ligand L with initial concentration [L_0_] and a set of proteins P_i_ with initial concentrations [P_0i_] and dissociation constants K_D,i_, the proportion of ligand bound to a specific protein P_k_ at equilibrium can be shown to be approximately

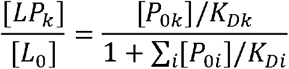

under the assumption that the free ligand concentration is much smaller than all dissociation constants. The denominator thus provides a tissue-specific promiscuity measure *π*_*k*_ that accounts for both binding affinity and local protein abundance. Since Ligand-Transformer predicts pK_D_ = -log_10_K_D_ = a, this can be rewritten as

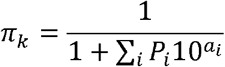

We also define the unweighted mean predicted binding affinity across the human proteome

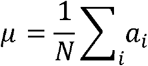

where N is the number of proteins in the reference set and ai is the predicted pK_D_ to protein i. We evaluated tissue-specific promiscuity across blood plasma, liver, brain, the HEK293 cell line, and a whole-body protein-abundance-weighted average using expression data from PaxDB^55^. All tissue-specific formulations correlated almost perfectly with the unweighted mean (Spearman ρ > 0.96 in every case; **Figure 1**), and we therefore adopted the unweighted mean as our primary promiscuity metric for all subsequent analyses.

### Specificity score

For each molecule, we computed the mean (μ) and standard deviation (σ) of its predicted binding affinities across all 15,405 proteins in the reference proteome. The specificity score for a given target protein T was defined as the Z-score

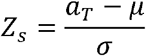

where *α*_*T*_ is the predicted pK_D_ of the molecule to target T. This score quantifies how many standard deviations the target affinity lies above the proteome-wide mean of the molecule, providing a normalised measure of selectivity that jointly accounts for target engagement and global off-target liability.

### Ligand-Transformer

Binding affinity predictions were generated using Ligand-Transformer, a deep-learning model that predicts pK_D_ values for small-molecule-protein pairs from their sequences and molecular structures. Ligand-Transformer encodes proteins using intermediate representations from AlphaFold-2^56^. Specifically, multiple sequence alignment (MSA) embeddings and pairwise amino acid feature embeddings capture evolutionary and geometric information. Ligands are encoded using GraphMVP^57^, a self-supervised graph neural network that derives molecular representations from SMILES strings with implicit knowledge of three-dimensional geometry. These protein and ligand representations are combined through a cross-modal attention module comprising twelve transformer-like layers, followed by separate affinity and distance prediction heads. A key advantage of Ligand-Transformer over structure-based docking methods is its ability to predict binding affinities for disordered proteins, which lack stable three-dimensional structures and are therefore inaccessible to conventional approaches.

### Datasets

A list of 19,843 proteins in the human proteome, along with their amino acid sequences, was downloaded from InterPro^58^. Because the computational cost of Ligand-Transformer inference scales quadratically with protein length, proteins longer than 730 amino acids (approximately 20% of the total) were excluded, yielding a final reference set of 15,405 proteins. This set spans the majority of human protein classes and interaction domains, and no excluded proteins featured among the top 10% most abundant proteins across any of the tissues investigated.

Protein abundance data across human tissues were obtained from the Protein Abundance Database (PaxDB)^55^, which provides whole-organism protein abundance averages covering approximately 85,000 proteins across twelve tissues in Homo sapiens.

Drug data were obtained from DrugBank 6.0^48^, from which 11,670 drugs with SMILES codes were extracted. Of these, 4,182 were labelled as approved, 2,663 were classified as small-molecule drugs, and 2,160 had at least one annotated protein target. Each drug was categorised by therapeutic function using the Description and Indication fields in DrugBank. The TargetMol compound library (https://www.targetmol.com/) provided 1,665,194 small molecules with SMILES codes, which served as the source of decoy compounds in enrichment analyses.

### Training OmniBind

OmniBind is built on the Chemprop architecture^42,43^, a PyTorch-based framework for training message-passing neural networks (MPNNs) for molecular property prediction. Training proceeded in two stages.

In the first stage, we generated ground-truth training labels by using Ligand-Transformer to predict the binding affinity of 61,677 molecules (11,670 from DrugBank and 50,007 from TargetMol) to all 15,405 proteins in the reference proteome. From these complete proteome binding profiles, we computed promiscuity scores (mean affinity) and standard deviations for each molecule. We refer to these as complete proteome promiscuity (CPP) scores. An initial MPNN trained directly on these 61,677 CPP labels achieved a best mean squared error (MSE) of only 0.41 after extensive hyperparameter optimisation, indicating insufficient training data. To augment the training set without incurring the full cost of proteome-wide Ligand-Transformer inference, we trained a random forest regressor to estimate promiscuity scores and standard deviations from a subsample of only 173 protein affinities (approximately 1% of the proteome). This intermediate model achieved an MSE of 0.03 against the CPP scores, and was used to generate 200,935 additional estimated promiscuity labels, termed representative protein promiscuity (RPP) scores, for TargetMol molecules at a fraction of the computational cost.

In the second stage, the final OmniBind MPNN was trained on the combined dataset of 61,677 CPP scores and 200,935 RPP scores. The model takes a SMILES string of a molecule as input and jointly predicts both the promiscuity score and the standard deviation of its binding profile. Data were split 80/10/10 into training, validation, and test partitions, and training was conducted for 100 epochs with hyperparameter optimisation. Final evaluation was performed on an additional held-out set of 1,000 TargetMol molecules with known CPP scores (**Figure S2**).

### Experimental promiscuity estimation

To validate OmniBind promiscuity scores against experimental data, we compiled small molecule-protein binding affinity data from three independent databases: ChEMBL^50^ (pKi and IC50 values), the Psychoactive Drug Screening Program (PDSP) Ki Database (pKi values), and the Safety Pharmacology Database (SPD)^45^ (AC50 values).

For the SPD, promiscuity was estimated for each molecule as the proportion of tested proteins with an associated AC_50_ ≤ 10 μM, as defined in the original study^45^. For ChEMBL and the PDSP, a common set of proteins with complete binding data across all molecules of interest was not available. We therefore identified the x proteins with the greatest number of associated binding measurements and retained only those k molecules for which data were available for at least a fraction i of these proteins. Parameters were chosen to balance protein coverage against the number of qualifying molecules: ChEMBL Ki (x = 5, i = 0.5, k = 1,247); ChEMBL IC_50_ (x = 5, i = 0.6, k = 177); PDSP Ki (x = 10, i = 0.6, k = 180); SPD AC_50_ (k = 893). For each qualifying molecule, a proxy promiscuity score was computed as the mean affinity across the available proteins.

### Maximum unbound plasma concentration (C_max_) data

Cmax measurements were obtained from the Safety Pharmacology Database (SPD)^45^ and the Pharmacokinetics Database (PKDB)^46^, filtered for blood plasma values from healthy adults. Where multiple C_max_ values existed for the same drug after filtering, the arithmetic mean was taken. To assess whether correlations between C_max_ and promiscuity were confounded by molecular size, we additionally computed correlations between C_max_ and log_10_(molecular weight) for the same drug sets.

### Affinity to best target and drug enrichment analysis

Drugs lacking an annotated protein target, with a target protein sequence exceeding 1,000 amino acids, or labelled as withdrawn were excluded, leaving 1,383 drugs for analysis. For each drug, we predicted its binding affinity to all annotated DrugBank targets using Ligand-Transformer and selected the highest-affinity target to obtain an affinity to best target (ABT) score. This was combined with the OmniBind promiscuity score and standard deviation to compute a specificity score for each drug-target pair.

To quantify the practical value of the specificity metric, we assessed its ability to distinguish approved drugs from random compounds. For each drug target, the corresponding drug was ranked alongside 1,000 randomly sampled TargetMol decoys, for which both Ligand-Transformer affinity predictions and OmniBind specificity scores were computed. Ranking performance was quantified using the Boltzmann-enhanced discrimination of the receiver operating characteristic (BEDROC) score^45^ at α = 10, 20, and 80, and enrichment factors at the 1%, 5%, 10%, and 20% thresholds. Drug categories for which specificity is pharmacologically less relevant (antiseptics, dermatologicals, diagnostics, nutritional supplements, and ophthalmics) were excluded from this analysis.

### Cross-validated enrichment analysis

To assess the robustness of the enrichment results, we performed a five-fold cross-validated analysis in which the decoy set was partitioned into five non-overlapping subsets of 192 molecules each. In each fold, drugs were ranked against the corresponding decoy subset and BEDROC and enrichment factor metrics were computed. The significance of the improvement from specificity over affinity-alone ranking was assessed using one-sided paired Wilcoxon signed-rank tests across folds.

### Generalisability across affinity predictors

To test whether the specificity improvement is robust to the choice of binding affinity model, we repeated the five-fold enrichment analysis using SSM-DTA, an independent drug–target affinity predictor, in place of Ligand-Transformer. As a further validation, we used Boltz-2 to predict the binding affinity of 63 approved drugs to each of the 53 proteins in the Eurofins SafetyScreen^46^ panel (https://www.eurofinsdiscovery.com/solution/safety-panels). We computed the mean Boltz-2 affinity score for each drug across the panel proteins and correlated these values with Ligand-Transformer-derived promiscuity scores. We additionally compared the frequency with which drugs classified as highly promiscuous (mean pK_D_ ≥ 5.85) appeared among the strongest 10% of Boltz-2 binders across the 53 panel proteins relative to low-promiscuity drugs, using a Mann-Whitney U test.

### OmniBind web server

The OmniBind web server accepts individual or batch SMILES submissions and returns promiscuity scores, binding profile standard deviations, and optional specificity scores. The server runs the trained MPNN model and returns predictions within seconds for typical batch sizes. Users may optionally supply a target protein identifier and a predicted or measured binding affinity to obtain a specificity score alongside the promiscuity prediction.

### Statistical analysis

Spearman’s rank correlation coefficients were used to assess monotonic associations between promiscuity measures and between computational and experimental promiscuity estimates. Pearson correlation coefficients were used for linear relationships, including promiscuity versus molecular weight and C_max_ analyses. Two-sided Student’s t-tests were used to compare mean promiscuity scores between approved drugs and random compounds. Mann-Whitney U tests were used for non-parametric comparisons of Boltz-2 binding frequencies between promiscuity classes. One-sided paired Wilcoxon signed-rank tests were used to assess significance in cross-validated enrichment comparisons. All p-values are two-sided unless otherwise specified. No corrections for multiple comparisons were applied, as all analyses were pre-specified.

### Data availability

Human proteome sequences were obtained from InterPro^58^ (https://www.ebi.ac.uk/interpro/). Drug annotations and SMILES codes were obtained from DrugBank 6.0^48^ (https://go.drugbank.com/). Protein abundance data are available from PaxDB^55^ (https://pax-db.org/). Experimental binding affinity data were obtained from ChEMBL (https://www.ebi.ac.uk/chembl/), the PDSP Ki Database (https://pdsp.unc.edu/databases/kidb.php)^51^, and the Safety Pharmacology Database. Pharmacokinetic data were obtained from the SPD^45^ and PKDB^46^ (https://pk-db.com/). The TargetMol compound library is available at https://www.targetmol.com/.

### Code availability

The OmniBind web server is available at [URL to be provided upon publication]. Source code for OmniBind, including trained model weights and analysis scripts, will be deposited in a public repository upon publication.

