## Supplementary figures for "OmniBind: Proteome-Wide Promiscuity Predictions for Early-Stage Drug Screening"

**SUPPLEMENTARY INFORMATION**

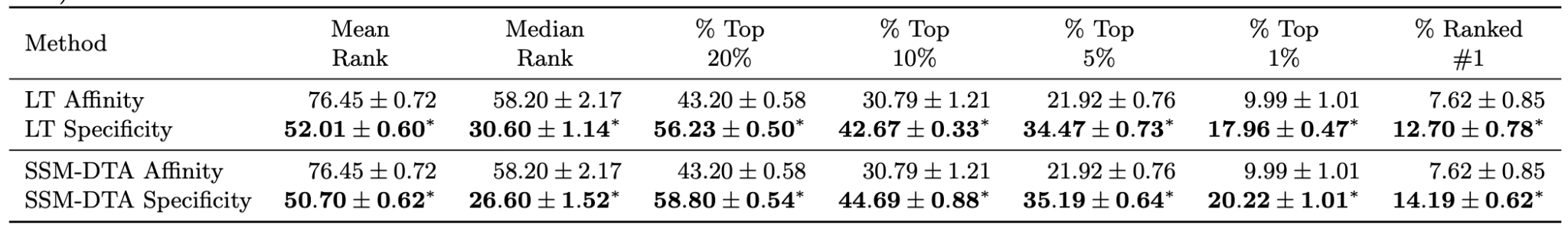

**Table S1.** **Specificity advantage is robust across cross-validation folds and affinity predictors.** Drug ranking performance across five-fold cross-validated decoy splits (n = 192 decoys per fold). FDA-approved drugs were ranked among TargetMol decoys using either predicted target affinity or the combined specificity score, with affinities computed by Ligand-Transformer (LT) or SSM-DTA. Columns report mean rank, median rank, and the percentage of drugs falling within the top 20%, 10%, 5%, and 1% of the ranked list, as well as the percentage ranked first overall. Values are mean ± standard deviation across folds. Asterisks denote metrics where specificity significantly outperformed affinity (one-sided paired Wilcoxon signed-rank test, p < 0.05).

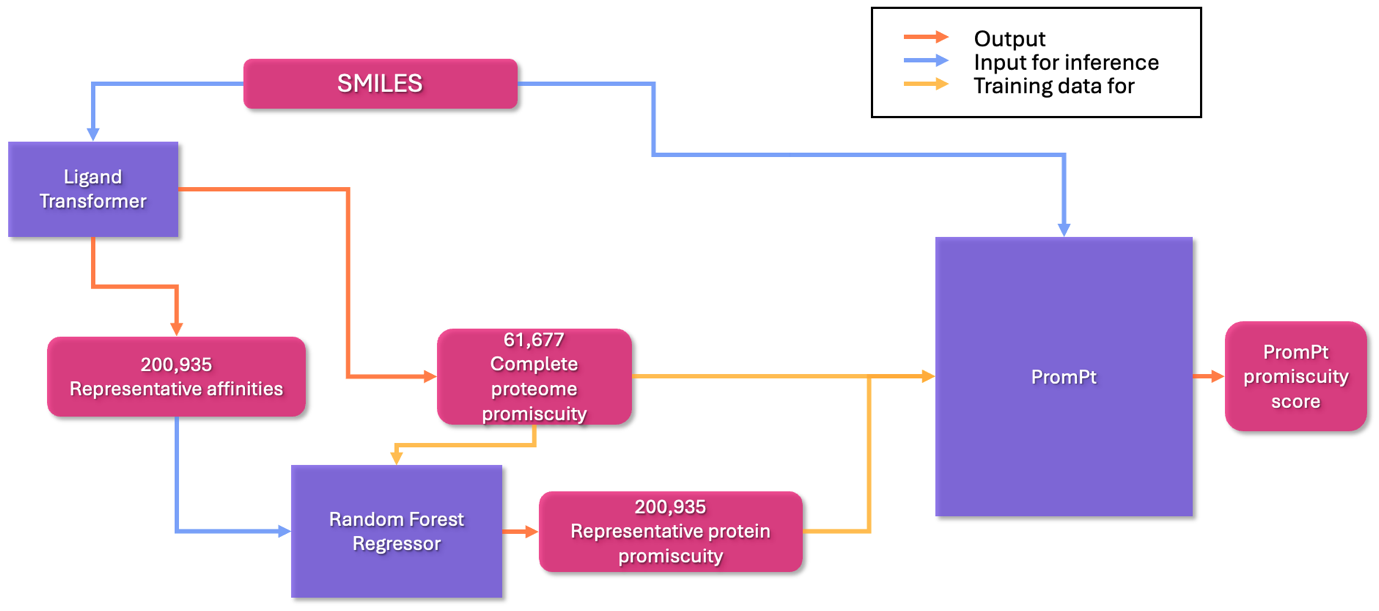

**Figure S1**. **Schematic overview of the OmniBind training pipeline.** Workflow used to construct the OmniBind promiscuity predictor. Ligand-Transformer was first used to compute full proteome-wide binding profiles for a set of small molecules across 15,405 human proteins. From these profiles, promiscuity scores (mean predicted affinity) and standard deviations were calculated. To expand the training set efficiently, a random forest model was trained to estimate promiscuity from affinities to a subset of representative proteins (~1% of the proteome), enabling rapid generation of additional training labels. These labels, together with the full proteome promiscuity scores, were then used to train the final OmniBind message-passing neural network (MPNN), which predicts promiscuity directly from a compound’s SMILES representation.

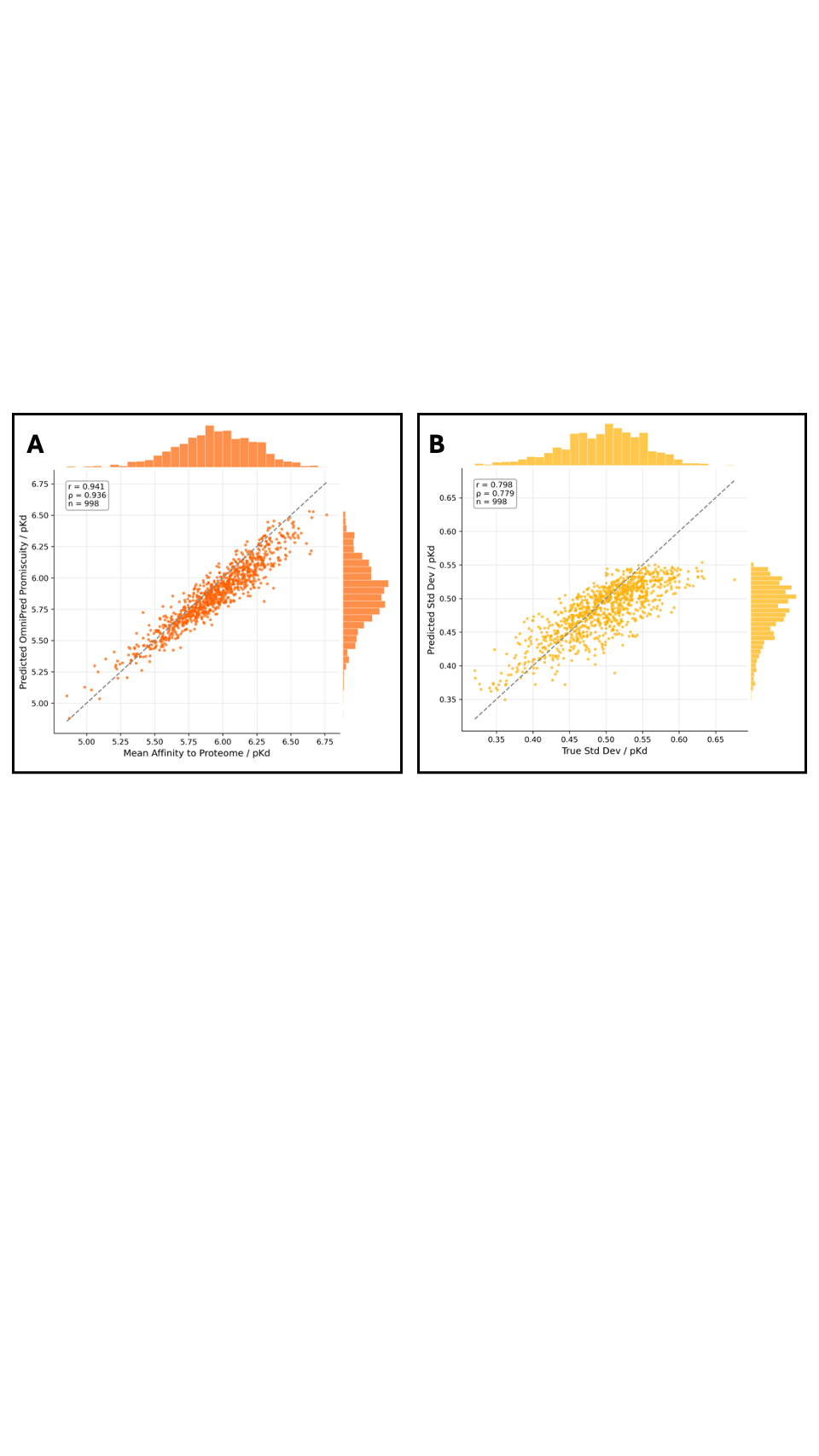

**Figure S2.** **Accuracy of OmniBind predictions of proteome-wide binding statistics.** Scatter plots comparing OmniBind predictions with reference values derived from Ligand-Transformer proteome-wide binding profiles for 1,000 held-out molecules. **(a)** Predicted versus reference mean binding affinity across 15,405 human proteins (promiscuity score). **(b)** Predicted versus reference standard deviation of binding affinities across the same proteome-wide distribution. Points correspond to individual compounds. OmniBind accurately reproduces both statistics of the binding profile with mean squared errors of 0.016 pK_D_ for the mean affinity and 0.001 for the standard deviation, demonstrating that the model captures the global binding tendency of small molecules across the proteome.

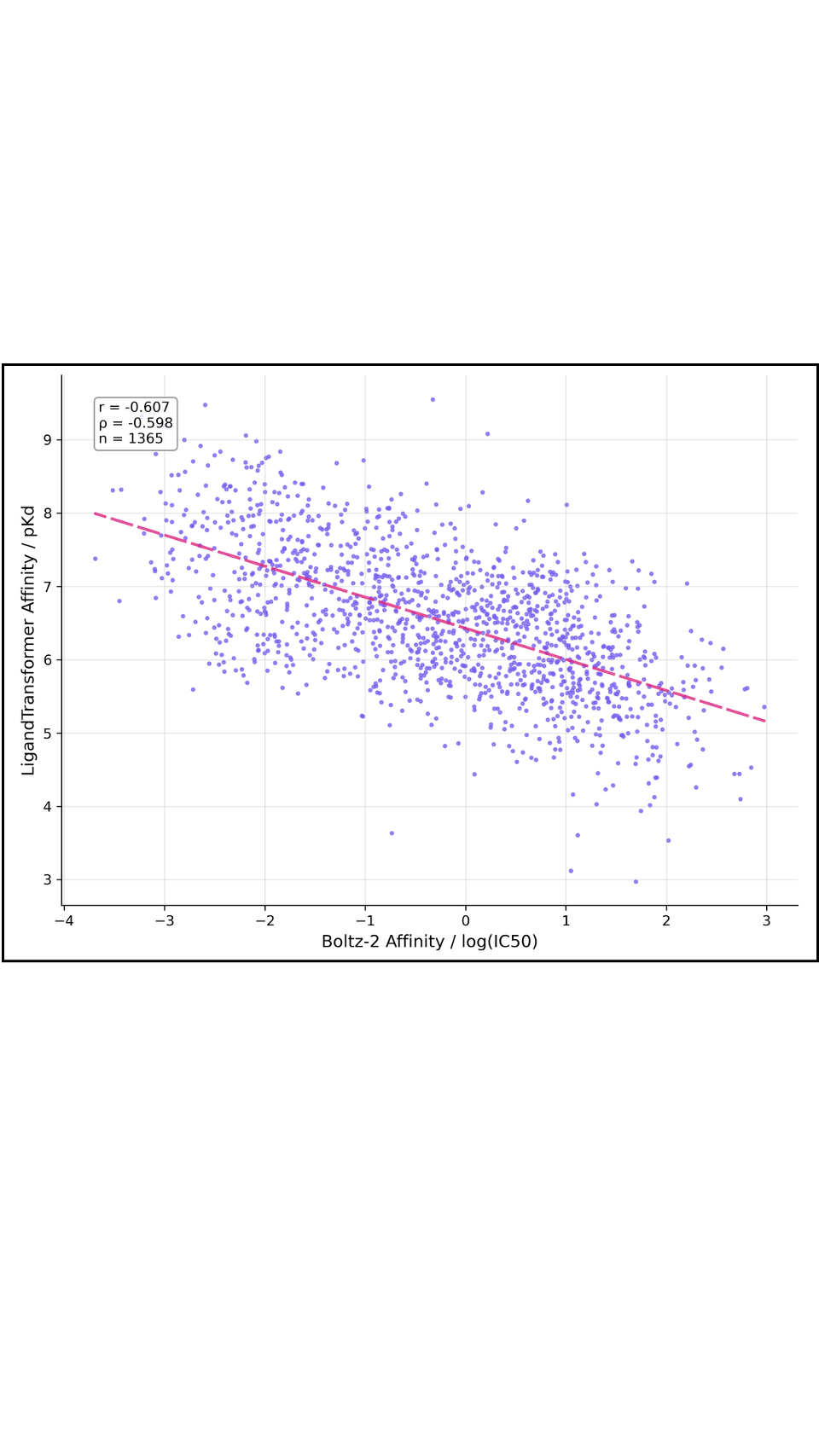

**Figure S3. Correlation between Ligand-Transformer and Boltz-2 affinity predictions.** Scatter plot comparing predicted binding affinities for drug–target pairs obtained with Ligand-Transformer and Boltz-2. Each point represents a compound–protein interaction. The two methods show strong agreement (|r| > 0.6, p < 0.001), indicating that the affinity predictions used to compute the promiscuity metric are broadly consistent with those produced by an independent structure-based predictor.

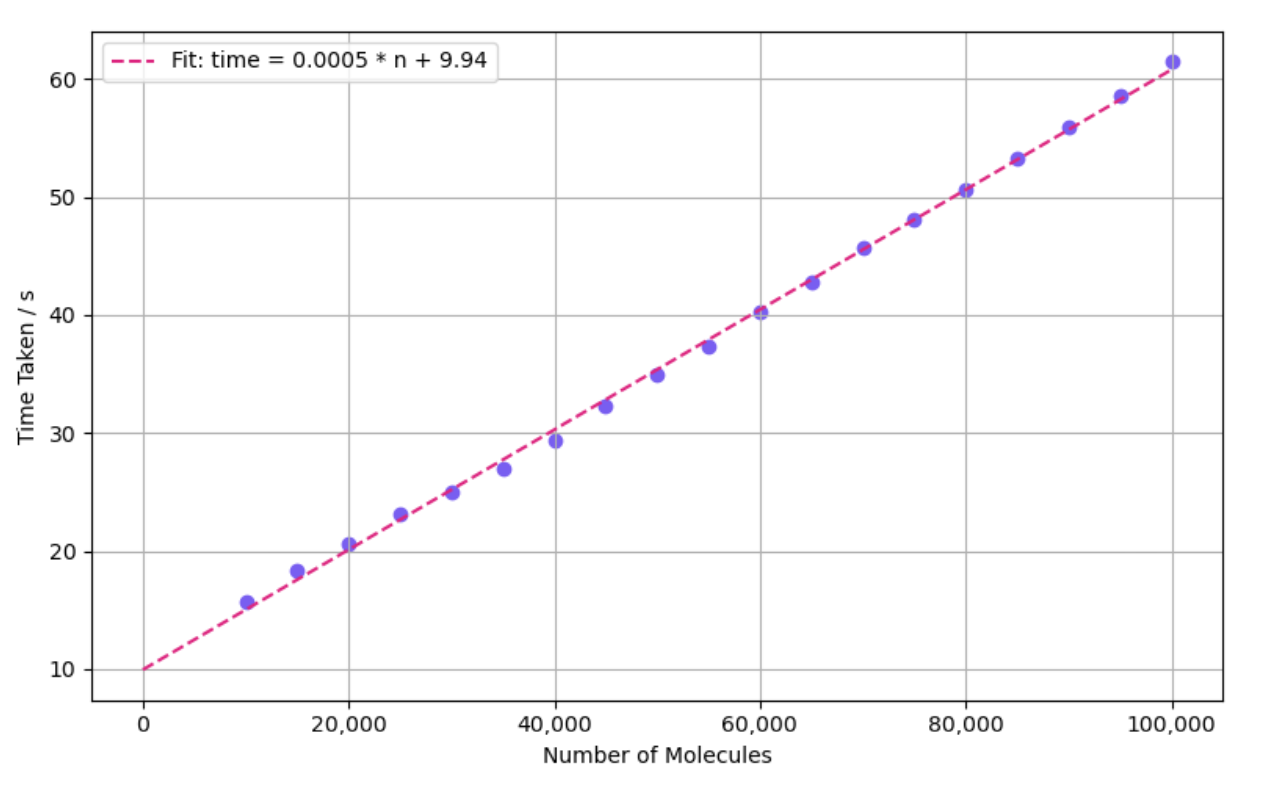

**Figure S4.** **Computational efficiency of OmniBind predictions.** Prediction time required for OmniBind as a function of the number of input molecules. The model exhibits approximately linear scaling with dataset size, enabling high-throughput promiscuity estimation. A regression line illustrates the relationship between runtime and number of molecules, demonstrating that OmniBind can process large compound libraries rapidly compared with full proteome profiling using Ligand-Transformer or other affinity predictors.

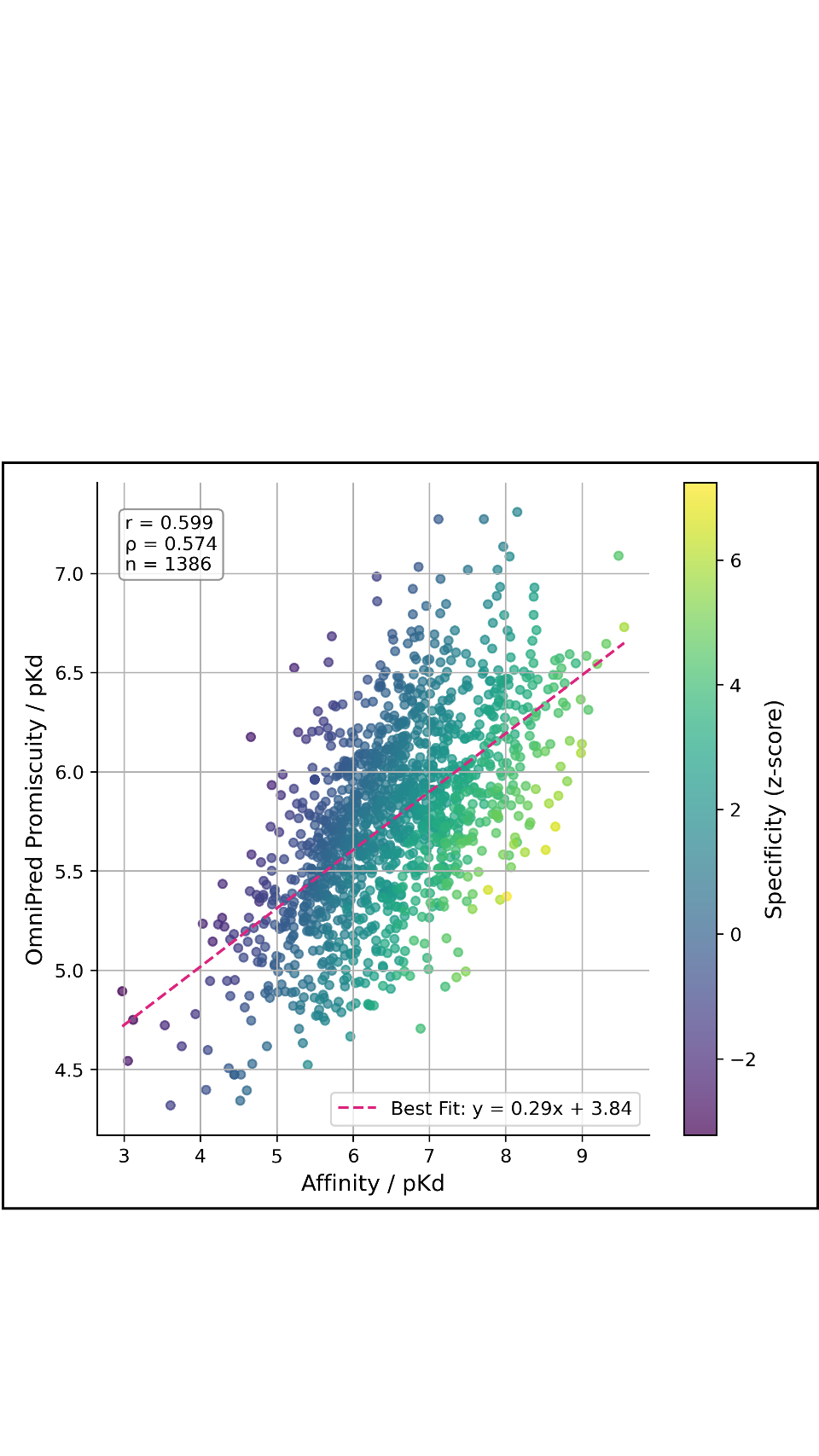

**Figure S5.** **Relationship between target affinity and proteome-wide promiscuity of approved drugs.** Scatter plot showing the correlation between the predicted affinity of each FDA-approved drug to its best target (affinity to best target, ABT) and its OmniBind promiscuity score, defined as the mean predicted binding affinity across 15,405 human proteins. Each point represents one drug. The positive correlation (r = 0.60, p < 0.001) indicates that compounds with stronger target affinity also tend to exhibit higher overall binding across the proteome, highlighting the importance of considering promiscuity alongside target affinity when evaluating drug candidates.

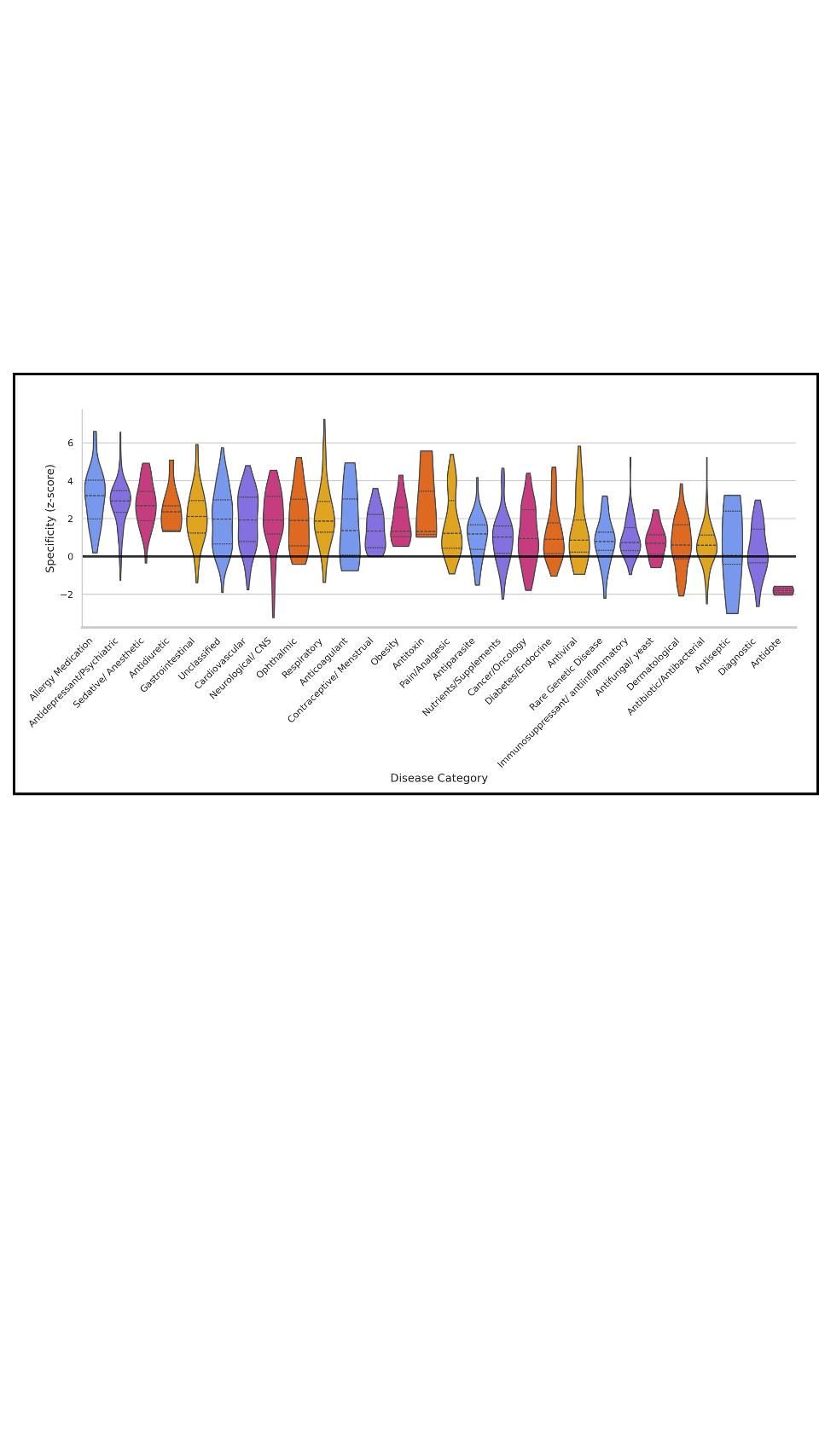

**Figure S6. Specificity scores of FDA-approved drugs across therapeutic categories.** Distribution of specificity scores for FDA-approved drugs grouped by therapeutic class. Specificity is defined as the Z-score of the affinity to the best target relative to a drug to the distribution of predicted affinities across the proteome, combining target affinity with proteome-wide promiscuity. Bars represent mean specificity scores within each category. Central nervous system-active drugs, including allergy medications, psychiatric drugs, and sedatives, show the highest specificity values, whereas categories such as diagnostics, antiseptics, and antibiotics exhibit lower specificity, consistent with the differing pharmacological requirements of these drug classes.

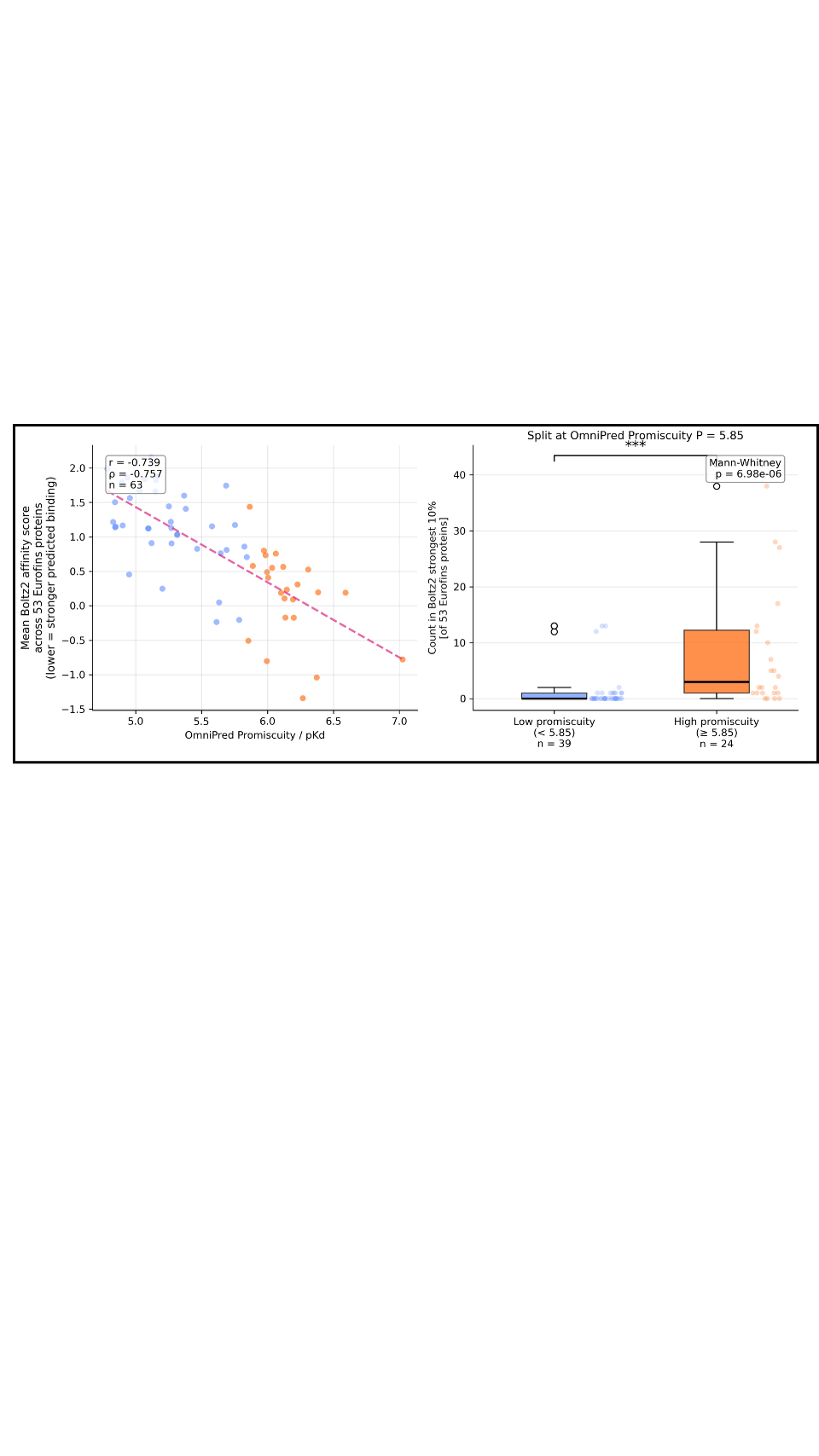

**Figure S7.** **Independent validation of promiscuity using Boltz-2 predictions on the SafetyScreen panel.** (a) Scatter plot showing the relationship between Ligand-Transformer–derived promiscuity scores and the mean binding affinity predicted by Boltz-2 across the 53 proteins in the Eurofins SafetyScreen panel for 63 FDA-approved drugs. Each point represents one drug; a strong correlation is observed (r=-0.74, p<10^-11^). (b) Enrichment of highly promiscuous drugs among the strongest predicted Boltz-2 binders. Drugs classified as highly promiscuous by Ligand-Transformer (mean pK_D_ ≥ 5.85) occur significantly more frequently within the top 10% of Boltz-2 binding scores across the panel than low-promiscuity drugs (Mann-Whitney p=7.0×10^-6^), supporting the consistency of the promiscuity metric across independent affinity predictors.
